# An opponent process for alcohol addiction based on changes in endocrine gland mass

**DOI:** 10.1101/2020.12.03.410365

**Authors:** Omer Karin, Moriya Raz, Uri Alon

**Affiliations:** Dept. Molecular Cell Biology, Weizmann Institute of Science, Rehovot Israel 76100

## Abstract

Consuming addictive drugs is often initially pleasurable, but escalating drug intake eventually recruits physiological “anti-reward” systems called opponent processes that cause tolerance and withdrawal symptoms. Opponent processes are fundamental for the addiction process, but their physiological basis is not fully characterized. Here, we propose an opponent processes mechanism centered on the endocrine stress-response, the HPA axis. We focus on alcohol addiction, where the HPA axis is activated and secretes β-endorphin, causing euphoria and analgesia. Using a mathematical model, we show that slow changes in HPA glands act as an opponent process for β-endorphin secretion. The model explains hormone dynamics in alcohol addiction, and experiments on alcohol preference in rodents. The opponent process is based on fold-change detection (FCD) where β-endorphin responses are relative rather than absolute; FCD confers vulnerability to addiction but has adaptive roles for learning. Our model suggests gland-mass changes as potential targets for intervention in addiction.

## Introduction

Drug addiction is a process in which the individual becomes increasingly occupied by drug-seeking and drug-taking behavior, escalates drug-taking over time, and has difficulties quitting. Addiction has several affective stages. While the contribution of each of these stages to the addiction process is debated (Wise and Koob, 2014), it is acknowledged that the stages are shared among many addictive drugs, including alcohol, opiates, and cocaine. The first stage is the initiation stage, which is related to positive reinforcement by drug administration. Next, the phenomenon of hedonic tolerance sets in: increasing amounts of drug are needed to produce the same effect. This leads to the maintenance stage in which the drug is taken in part to avoid the negative effects of withdrawal. Changes in learning and memory systems result in compulsive drug-seeking habits and increase subsequent risk of relapse (Hyman et al., 2006; Milton and Everitt, 2012; Robbins, 2002). Once drug-use is stopped, the withdrawal stage occurs, characterized by persistent negative affect (Koob, 2011; Koob and Le Moal, 2001). This provides negative reinforcement against cessation of drug use. Withdrawal gradually gives way to recovery (Figure 1A).

**Figure 1.**
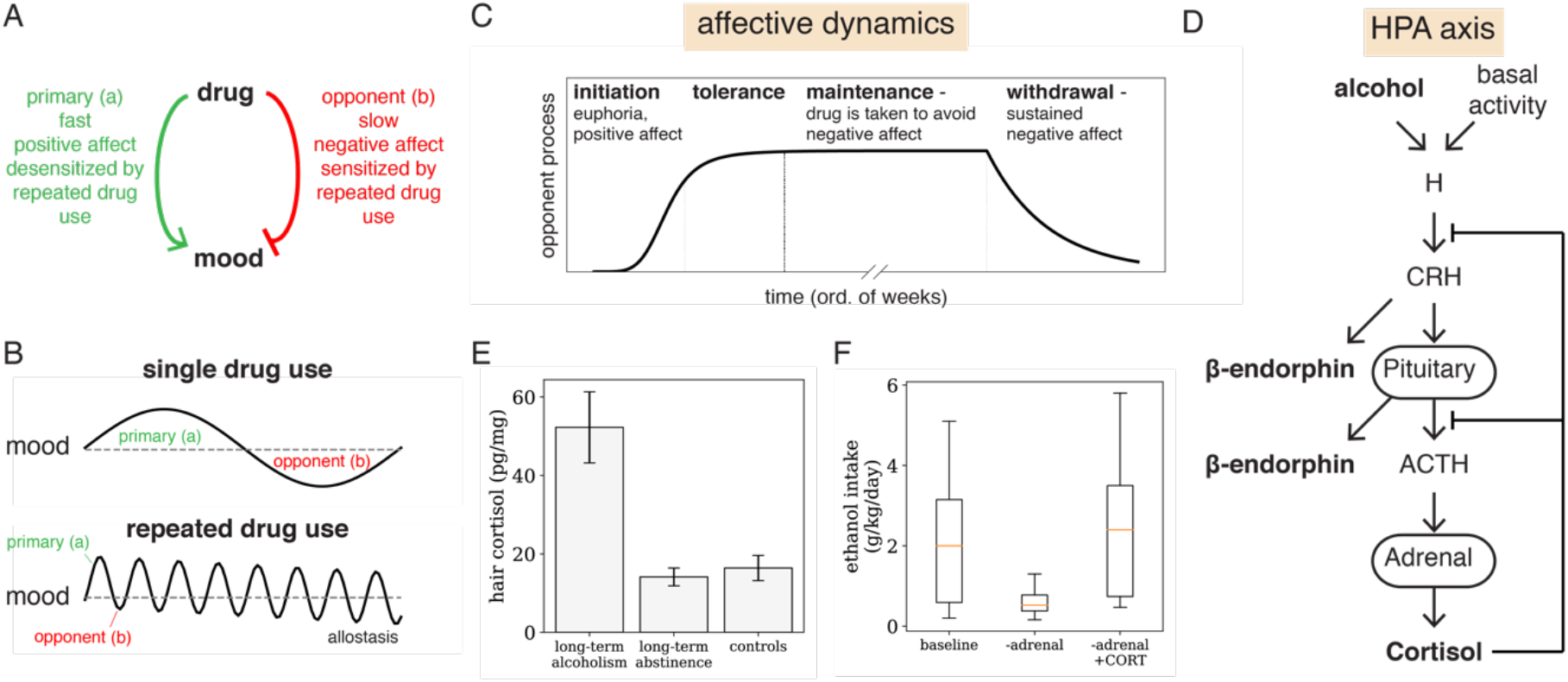
Opponent process theory explains the affective stages of addiction. (A) According to the opponent process theory of drug addiction (Koob and Le Moal, 2001; Koob and Volkow, 2010), the intake of an addictive drug activates two processes: a fast, primary process that causes the pleasurable effects of the drug, and a slow “anti-reward” opponent process which results in negative affect. Repeated intake over long periods of time desensitizes the primary process and sensitizes the opponent process, resulting in allostasis of mood and sustained negative affect after drug withdrawal. (B) Schematic illustrations of the effects of single and repeated drug intake on mood. (C) The opponent process framework can explain the stages of addiction to various drugs: initiation (weeks), tolerance that leads to maintenance (weeks to decades), and withdrawal (weeks to months). (D) Regulation of β-endorphin by alcohol. β-endorphin is secreted from POMC-neurons in response CRH from the hypothalamus, denoted H. CRH also causes pituitary corticotrophs to secrete ACTH, which is cleaved from the same peptide as β-endorphin. ACTH causes the adrenal gland to secrete cortisol, which, in turn, inhibits the secretion of CRH and the pituitary secretion of ACTH and β-endorphin. Cortisol is therefore is a candidate component of an opponent process mechanism for the rewarding effects of β-endorphin. This is supported by data indicating that cortisol is elevated after long-term alcoholism (E, from (Stalder et al., 2010)), and that alcohol preference is reduced in adrenalectomized animals and restored by cortisol replacement (F, data from (Fahlke et al., 1994)).

These affective changes have been studied by the opponent-process theory of addiction (Koob and Le Moal, 1997; Koob and Volkow, 2016; Solomon, 1980) (Figure 1ABC). This theory posits that chronic drug use activates anti-reward processes called opponent processes. Opponent processes are secondary slow processes that are activated by the drug and which antagonize the primary, pleasurable process (Fig 1A). After initial drug use, the opponent process causes a slight and slow-to-decay undershoot in mood. Repeated, extensive drug use sensitizes the secondary process and desensitizes the primary process (a phenomena known as “reward allostasis”), leading to deficient primary hedonic responses and exaggerated negative affect upon drug withdrawal. This explains the transition from the initial euphoric stage of drug use to the maintenance stage, where the drug is taken to avoid the negative effects of withdrawal that are due to the opponent process.

An important challenge for addiction research is to identify the molecular mechanisms of these opponent processes. The difficulty lies in the complexity of the physiological circuits that underlie addiction. These reward-processing circuits control pleasure, pain and reinforcement (Berridge and Robinson, 2016; Kelley and Berridge, 2002; Koob and Le Moal, 1997; Koob and Volkow, 2016; Solomon, 1980). Addictive drugs including alcohol, cocaine, and opiates affect the levels of specific neurotransmitters and other secreted molecules, which, in turn, affect the brain regions involved in reward processing (Koob and Volkow, 2016). Since these circuits are intricate and involve feedback over multiple timescales, mathematical models are essential for analyzing their relation with addiction dynamics and for pointing to potential opponent processes.

Most proposals for opponent processes involve neurotransmitter circuits, primarily dopamine, a key neuromodulator of learning and motivation (Berridge and Robinson, 1998; Glimcher, 2011; Schultz, 1998; Schultz et al., 1997; Wise, 2004). One such model was developed by Gutkin et al. to study nicotine addiction (Gutkin et al., 2006). Gutkin et al. analyzed the interaction between nicotine and nicotine receptors expressed in neurons which secrete dopamine. They proposed a model where on the fast time scale nicotine causes dopamine to be secreted and upregulates the phasic dopamine response, while on a much slower timescale chronic nicotine administration activates an opponent process which downregulates tonic dopamine responses. Upon nicotine withdrawal, this slow process causes an undershoot in tonic dopamine. Other proposed opponent processes include the recruitment of stress-associated neurotransmitters such as dynorphin and CRH in the amygdala, and various other neurochemical systems (reviewed in (Koob and Volkow, 2016)).

Since opponent processes are important for understanding addiction, it is useful to propose additional physiological systems which can provide opponent process properties.

Here we propose a new class of opponent processes, based not on neurotransmitters but instead on endocrine circuits. For this purpose, we model the dynamics of an endocrine system which plays an important role in alcohol addiction: the secretion of endogenous opioids following HPA axis activation (Figure 1D). Endogenous opioids control subjective reward, euphoria and pain sensitivity (Darcq and Kieffer, 2018; Drews and Zimmer, 2010; Gerrits et al., 2003; Gianoulakis, 2004; Kiefer et al., 2002; Kuzmin et al., 1997; Mitchell et al., 2012; Roberts et al., 2000; Roth-Deri et al., 2008; Trigo et al., 2009; Van Ree, 1996). They are secreted in response to stress after activation of the HPA axis (Nikolarakis et al., 1986; Tsigos and Chrousos, 2002), and affect dopamine release (Spanagel et al., 1992). In particular, β-endorphin, which causes euphoria and analgesia by binding to the same receptor as morphine, has been tightly linked with addiction to alcohol and other substances which activate the HPA axis (Gianoulakis, 2004; Kiefer et al., 2002; Roth-Deri et al., 2008; Trigo et al., 2009). Rodents that lack β-endorphin or its receptor (mu-opioid receptor) show diminished alcohol self-administration (Hall et al., 2001; Racz et al., 2008; Roberts et al., 2000), and deficiency in β-endorphin during the weeks after alcohol withdrawal is associated with anxiety in patients (Kiefer et al., 2002).

The importance of β-endorphin for alcohol addiction raises the question of what its opponent process may be. One candidate is the secretion of the stress hormone cortisol from the adrenal gland. Cortisol inhibits β-endorphin secretion both directly and by inhibition of CRH secretion. This was shown by experiments employing adrenalectomy, as well as by pharmacological manipulations such as the administration of dexamethasone and of the glucocorticoid synthesis inhibitor metyrapone (Guillemin et al., 1977; Nakao et al., 1978; Holaday et al., 1979; Lim et al., 1982; Rivier et al., 1982; Kreek et al., 1984; Hargreaves et al., 1987; Young, 1989). Cortisol is greatly elevated during periods of excessive alcohol consumption (Figure 1E, (Esel et al., 2001; Stalder et al., 2010)), and returns to baseline over a few weeks after alcohol withdrawal (von Bardeleben et al., 1989; Esel et al., 2001; Kiefer et al., 2002; Marchesi et al., 1997). Abolishing cortisol secretion by the removal of the adrenal glands diminishes alcohol preference in rodents. Alcohol preference in adrenalectomized animals is rescued by corticosterone replacement (Figure 1F) (Fahlke, 2000; Fahlke et al., 1994; Hansen et al., 1995; Lamblin and De Witte, 1996). Similar results were reported for cocaine (Goeders, 2002; Goeders and Guerin, 1996)). Since adrenalectomy removes the negative feedback from β-endorphin secretion (Young, 1989), this raises the possibility that cortisol secretion is an opponent process for β-endorphin. However, cortisol has a half-life of only 1 hour, similar to that of β-endorphin (Foley et al., 1979; McKay and Cidlowski, 2003). This is much faster than the timescale of days-weeks expected for an opponent process, suggesting that adrenal secretion of cortisol cannot by itself be the opponent process for β-endorphin secretion.

Here we propose that the opponent process for β-endorphin secretion may be due to structural changes of the HPA glands following repeated alcohol intake, namely the growth of the total mass of the hormone-secreting cells in the adrenal and pituitary glands. We show this by analyzing the dynamics of β-endorphin secretion following acute and repeated alcohol intake, using a recently-developed mathematical model of the HPA axis (Karin et al., 2020). The important aspect of this model is that it includes the changes in the total cell mass of the HPA glands due to cell proliferation and growth. This growth is caused by the HPA hormones, which act as growth factors for specific HPA glands. Changes in gland masses are well documented in conditions associated with HPA activation in humans (Amsterdam et al., 1987; Carey et al., 1984; Doppman et al., 1988; Ludescher et al., 2008; Nemeroff et al., 1992; Rubin et al., 1995, 1996), including enlarged adrenal glands in chronic alcohol abuse (Carsin-Vu et al., 2016). Adrenal weight in rodents also increases after alcohol administration (Adams and Hirst, 1984; Ɖikić et al., 2011; Mendelson et al., 1971).

The gland-growth mechanism adds a timescale of weeks to the hours-timescale of the HPA hormones. The model shows that while alcohol initially increases β-endorphin levels, persistent HPA activation by alcohol causes the HPA glands to grow, causing increased cortisol secretion which renormalizes β-endorphin levels and suppresses them upon withdrawal. *The gland masses thus provide an opponent process*, explaining data on β-endorphin and HPA hormones during alcohol addiction and withdrawal, as well as the effect of adrenalectomy and cortisol replacement on alcohol preference in rodents. The main message of this study is thus that a weeks-scale feedback loop in which HPA gland masses changes over time provides an opponent process.

Using the model, we analyze the fundamental reason for the HPA opponent process: fold-change detection (FCD) for β-endorphin dynamics in response to drug inputs. FCD is a property of biological circuits (Adler and Alon, 2018; Shoval et al., 2010) where the output of the circuit responds to *relative* changes in the input, rather than absolute changes. The model makes several predictions that can be tested experimentally. We show, using a minimal model of reward optimization, that FCD circuits that control subjective reward are uniquely fragile to addiction, because they maintain sensitivity to reward at increasing levels of drug intake.

In addition to this fragility, FCD control of subjective reward has two beneficial functions for reward-based learning. The first is the ability to learn across many orders of magnitude of rewards. The second benefit is that FCD implements an important concept from reinforcement learning, called potential-based reward shaping (Ng et al., 1999), which helps an individual to learn by adding auxiliary rewards that guide exploration. Thus, FCD has beneficial properties for learning, while it has fragility to addiction.

## Results

### A model for drug-induced regulation of β-endorphin

To characterize the HPA opponent process, we model the regulation of β-endorphin (Figure 1D). β-endorphin is an endogenous opioid that is secreted in response to hypothalamic CRH from corticotroph cells in the pituitary gland and from POMC neurons. Alcohol taking causes CRH secretion and hence β-endorphin secretion (Čupić et al., 2017; Lee et al., 2001, 2004; Rivier, 1996; Rivier et al., 1984; de Waele and Gianoulakis, 1993). In addition to causing the secretion of β-endorphin, CRH also activates the HPA endocrine cascade which results in the secretion of the stress hormone cortisol. Cortisol, in turn, provides negative feedback on CRH and β-endorphin. This feedback acts on the timescale of minutes to hours, and cannot by itself provide slow opponent properties on the timescale of days-weeks.

To address the slow timescale, we recently developed a mathematical model of the HPA axis on the time scale of weeks (Karin et al., 2020). We showed that these dynamics can explain the week-long changes in responses to CRH tests after withdrawal from prolonged alcohol use (Karin et al., 2020). Here we add β-endorphin to this model, and use it to study addiction.

The weeks-timescale in the model is due to growth of the total mass of the pituitary corticotroph cells that secrete ACTH and β-endorphin, and the adrenal cortex cells that secrete cortisol. Hereafter, we call such total cell masses “gland masses” for brevity, although the glands also contain other cell types. The mass of the gland affects the amount of hormone it secretes for a given input signal: twice the mass is assumed to result in twice the hormone secretion.

These cells constantly turn over, with a turnover time of weeks. Their major growth factors are the HPA hormones themselves: CRH causes corticotroph mass to grow (Gertz et al., 1987; Westlund et al., 1985; Carey et al., 1984; Horvath, 1988; Schteingart et al., 1986; O’Brien et al., 1992; Asa et al., 1992; Bruhn et al., 1984; Young and Akil, 1985; Gulyas et al., 1991; McNicol et al., 1988), and ACTH causes the adrenal cortex mass to grow (Dallman, 1984; Lotfi and de Mendonca, 2016; Swann, 1940; Ulrich-Lai et al., 2006).

To describe the dynamics of β-endorphin following alcohol intake, we added equations as described in (Methods, Figure 2A). In the model, alcohol-taking is described as an input *u_D_* that is additive with the basal circuit input *u_B_*, so the total input is *u* = *u_B_* + *u_D_* (Figure 2A). Persistent HPA activation by alcohol increases the hormone levels, and since these hormones act as growth factors for the glands, the gland masses grow on the timescale of weeks.

**Figure 2.**
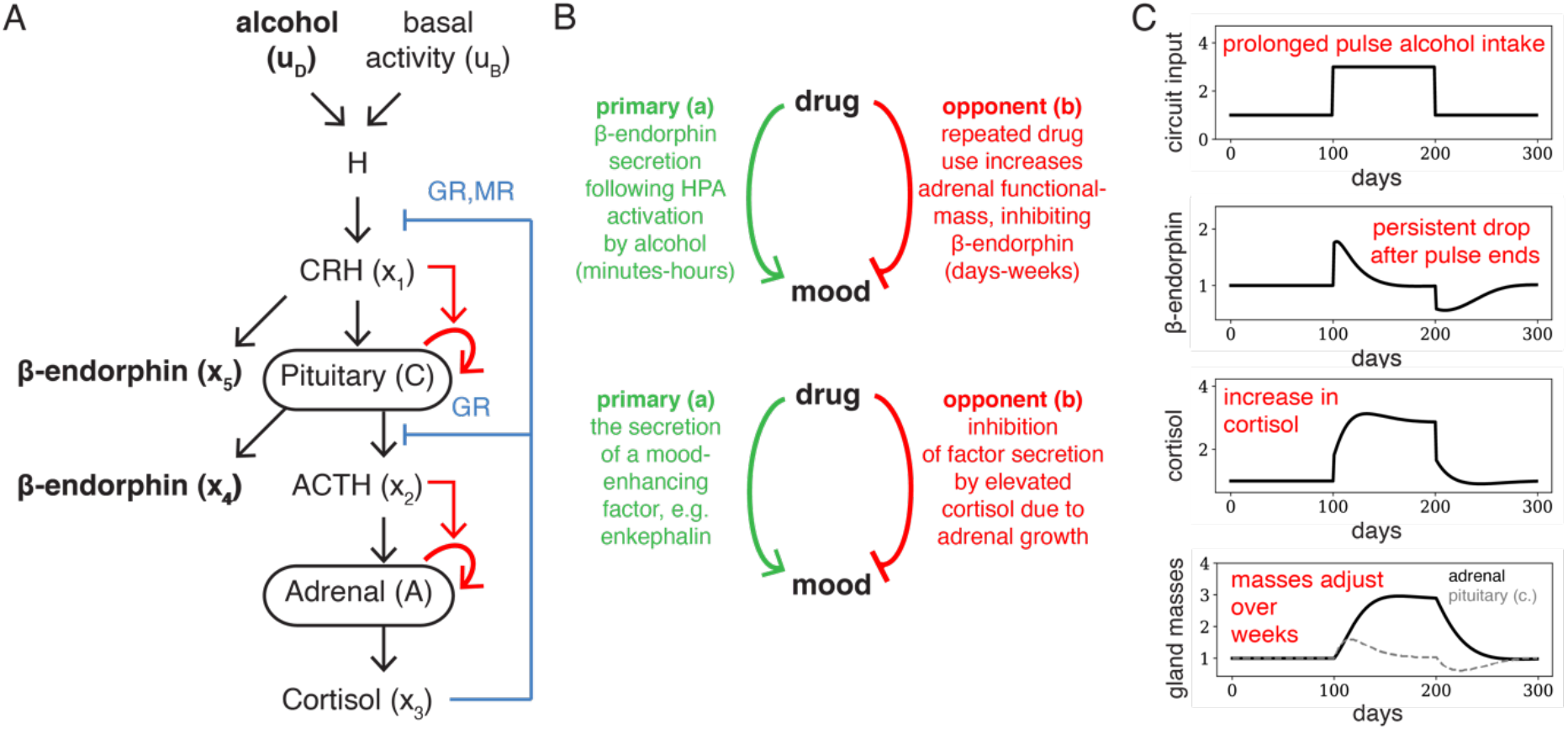
Opponent process for β-endorphin based gland-mass changes in the HPA axis. (A) We model HPA control of β-endorphin by adding β-endorphin to our recent model of the HPA axis (Karin et al., 2020) that incorporated week-timescale interactions for the gland masses (red arrows). CRH causes the growth of pituitary corticotroph cells, and ACTH causes the growth of the adrenal cortex cells. The model equations and parameters are provided in Methods. (B) The model provides opponent process properties, where the primary process is the secretion of β-endorphin following HPA activation, and the opponent process is changes in cortisol secretion due to change in adrenal mass following prolonged HPA activation. The results generalize to other mood-enhancing factors that are secreted in response to alcohol intake, and that are inhibited by cortisol. (C) A prolonged increase in alcohol-taking causes β-endorphin to initially increase (primary process) but then return to baseline (opponent process). The reason for this is the growth of the gland masses, which increases the negative feedback of cortisol on β-endorphin. The return of β-endorphin to baseline develops over the timescale of weeks, which is much slower than the half-life of β-endorphin. Stopping alcohol intake after months leads to a drop in β-endorphin levels which lasts for a few weeks, together with an undershoot in pituitary corticotroph mass. Shown is pituitary β-endorphin; similar secretion patterns occur for β-endorphin secreted from POMC neurons.

To test whether the glands can implement an opponent process for β-endorphin secretion, we consider a months-long input pulse that corresponds to a prolonged increase in average alcohol-taking (Figure 2BC, similar results are obtained by simulating discrete drinking episodes). Initially, β-endorphin and cortisol levels increase, because the drug activates the HPA axis. The increase in β-endorphin after drug intake is the ‘a-process’ in the opponent-process model, and has a timescale of a few hours. This occurs without significant change in the gland masses.

Over the next several weeks, glands masses adjust, as described in Karin et al. (Karin et al., 2020). A change in gland mass is assumed to cause a proportional change in the rate of secretion of the hormone produced by the gland. This is because we consider the secretion rate of each hormone as a product of three factors: the effect of the signals (agonists like CRH and antagonists like glucocorticoids), the maximal secretion capacity per unit biomass, and the total biomass of the cells. These factors are separated by timescales: signals can change over minutes, maximal secretion capacity over hours (due to protein expression changes), and total cell biomass over days-weeks (due to hypertrophy and hyperplasia). Thus, at a given level of signals and at a given maximal secretion capacity per unit biomass, the production of the hormone is proportional to the cell total biomass: doubling the biomass doubles secretion. The model takes into account both the fast changes in signals and the slow changes in gland masses. The separation into these factors allows us to consider the effects over weeks of gland mass changes and disentangle them from faster effects. It also allows us to follow changes in the mass of two cell types-pituitary corticotrophs and adrenal cortex cells - which change at the same time.

With these considerations, we see that HPA activation causes growth of pituitary corticotroph mass over weeks, due to the action of CRH as a growth factor. The enlarged biomass enhances β-endorphin secretion. In parallel, HPA activation makes the adrenal mass grow because of increased levels of its growth factor, ACTH. Growth of the adrenal mass causes increased secretion of glucocorticoids, which eventually suppresses β-endorphin secretion. This acts to renormalize β-endorphin levels back to baseline (mathematically, this is due to the integral feedback in Eq. 4–5 in Methods). The change in the adrenal mass is the slow ‘b-process’ in the opponent process model. It causes suppression of beta-endorphin on a timescale of days-weeks.

The same effects cause an undershoot of β-endorphin which lasts for weeks-months upon alcohol withdrawal after prolonged intake. The undershoot is caused by the enlarged HPA gland masses, which return to their baseline size over several weeks. These dynamics are qualitatively similar to those predicted by Gutkin et al. for the effect of nicotine on tonic dopamine through nicotinic receptors (Gutkin et al., 2006), but arise due to different physiological processes.

These opponent-process stages of gland growth and shrinkage occur regardless of model parameters, as long as the turnover time of the gland cells is much slower than the halflives of the hormones. The opponent process is thus a robust prediction of the model.

The model explains several important observations regarding alcohol addiction. It explains the large increase in average cortisol levels seen after prolonged drinking (Stalder et al., 2010), the acute elevation of β-endorphin levels after alcohol administration (Frias et al., 2002; Marinelli et al., 2003), and the prolonged drop seen in β-endorphin levels after alcohol withdrawal (Aguirre et al., 1990; Esel et al., 2001; Inder et al., 1995; Kiefer et al., 2002; Marchesi et al., 1997; Vescovi et al., 1992). More generally, it captures the dynamics of dysregulation of HPA hormones during alcohol withdrawal (von Bardeleben et al., 1989; Karin et al., 2020). These effects are due in the model to the slow changes in the gland masses.

### Opponent-process behavior is due to a fold-change detection property

The opponent-process behavior of β-endorphin is due to a systems-level feature of the HPA model called fold-change detection (FCD, Figure 3A). This property has been extensively studied in systems biology. A system with FCD has an output whose entire dynamics depends only on *relative* changes in input, rather than absolute changes (Adler and Alon, 2018; Shoval et al., 2010). All systems with FCD also have the property of exact adaptation, where the output adapts precisely back to baseline after step changes in its input (Ferrell, 2016; Shoval et al., 2010).

**Figure 3.**
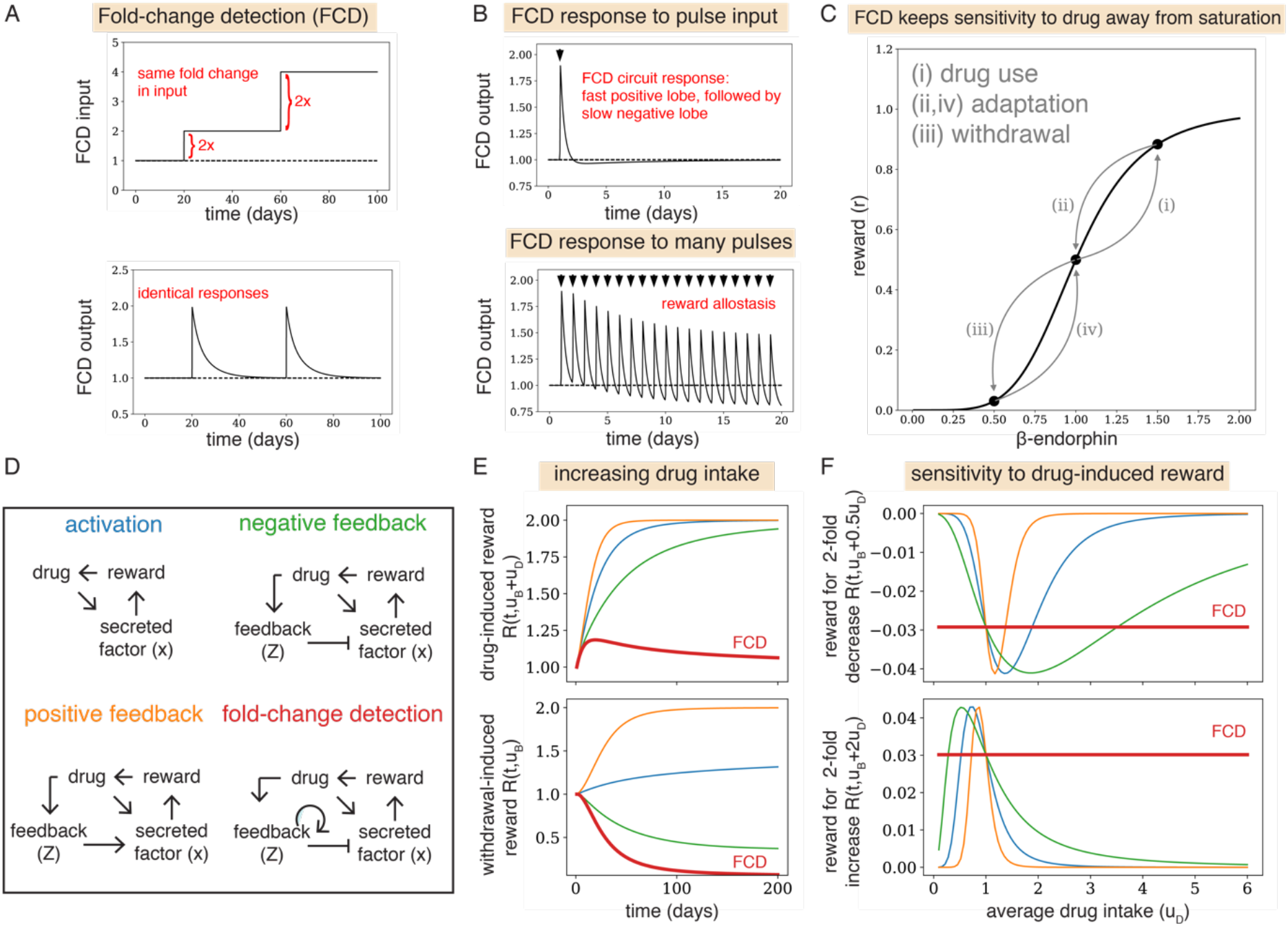
FCD control of subjective reward provides the hallmarks of the opponent process model and is prone to addiction. (A) Foldchange detection is a property of biological circuits where, after adaptation, the circuit responds to relative rather than absolute changes in input. (B) A pulse input (dark arrow) causes a doubled lobed response with a fast-positive response followed by a slow-negative response, so the overall integral is zero. This property entails that positive hedonic responses (“a-process”) are balanced by subsequent negative hedonic responses (“b-process”). Repeated pulses desensitize the positive response and sensitize the negative response so the average overall output level is constant, explaining reward allostasis. (C) FCD circuits maintain sensitivity to drug intake by keeping the system away from saturation. We demonstrate this with a schematic illustration, using a generic hill-function which maps between β-endorphin levels and subjective reward 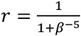. Initially, the individual will increase drug intake to maximize subjective reward. This causes β-endorphin to increase, so that subjective reward rises closer to saturation (arrow marked (i)). Then, due to FCD, β-endorphin secretion adapts back to baseline over weeks, and subjective reward returns to baseline (ii). The circuit maintains its sensitivity relative to the new baseline, so that a larger drug increase is required to get the same rise in subjective reward. Finally, when the individual ceases to consume the drug, β-endorphin and subjective reward fall (iii), until β-endorphin re-adapts and returns to baseline over weeks (iv). (D) Four prototypical circuits for the control of subjective reward, FCD (red lines) where subjective reward depends on the fold-change of drug input, negative or positive feedback (green and orange lines, respectively), where a slow process Z feeds back on subjective reward, and activation (blue lines), where the drug input translates directly into subjective reward (equations in Methods). (E) Simulations of drug intake that rises linearly with time, *u*(*t*) = 1 + 0.015*t*. In all circuits except FCD, drug-induced reward rises with the increase in drug intake. Withdrawal mood drops without bound only for the FCD circuit. (F) The FCD circuit (red line) maintains preference for an increase in drug intake even at high levels of drug intake, whereas other circuits show diminishing preference at high intake levels.

The HPA model shows that the dynamic response of CRH depends only on relative changes in input, rather than absolute changes (see Methods for the proof). Because β-endorphin is secreted in response to CRH, it also has the FCD property (see Methods for the proof both for pituitary and brain secretion of β-endorphin). The FCD property holds at low to moderate levels of HPA-axis activation, when the low-affinity GR activation is weak. At higher levels of HPA activation, GR feedback ameliorates FCD and causes reduced responses to the same fold-changes in input.

The reason for FCD is that gland masses grow in response to an increase in input on the timescale of weeks. The increased gland mass results in higher levels of cortisol. Cortisol levels thus rise proportionally to the input level. Since cortisol inhibits CRH and β-endorphin secretion, the latter are “normalized” by the input and show exact adaptation back to their baseline (Figure 2B), as well as fold-change responses (Figure 3A).

The FCD property provides the essential hallmarks of an opponent process (Figure 3B), as defined in (Koob and Volkow, 2010). These hallmarks concern the hedonic responses to single and repeated episodes of drug intake. After drug intake, a positive hedonic response occurs, and matches the intake’s duration and intensity. The negative hedonic response, which is due to the opponent process, follows the positive response, with a slow and prolonged decay. Repeated exposure to the drug reduces the positive response and increases the negative response, so the overall average hedonic response remains constant (this is referred to as reward allostasis).

All of these hallmarks are captured by FCD (Figure 3B). The positive response of an FCD circuit to a pulse input is always followed by a slow trailing negative response, because the overall integral of the response must be zero (Figure 3B, proof in Methods). Because FCD circuits adapt to the average input level, repeated exposure to drug intake desensitizes the positive response and sensitizes the negative response, leading to reward allostasis (Figure 3B). We therefore propose that FCD, a concept from systems biology that is common in sensory circuits, aligns with the essential features of the opponent process theory.

To make these notions more precise, we attempted to connect these FCD properties to the reward aspects of the opponent process theory more formally. For this purpose, we need to operationally define three variables in quantitative terms: subjective reward, drug reward, and withdrawal mood. We denote by *u_B_* the basal input to the HPA axis and by *u_D_*(*t*) the input due to drug intake at time *t*, so the total input to the HPA axis is *u*(*t*) = *u_B_* + *u_D_*(*t*). We take the generic assumption that the effect of β-endorphin on subjective reward is a saturating (Hill) function, which describes the saturation of the opioid receptors, or downstream saturation of subjective reward in the brain. Thus, the instantaneous subjective reward is 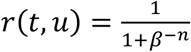 where *β* is the level of β-endorphin and u is the HPA axis input is.

The total subjective reward *R* from taking a drug can be calculated by the accumulated subjective reward over a time period after taking the drug. As customary in the reinforcement learning literature, total subjective reward is given by the discounted integral over the instantaneous reward: 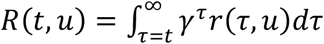 where *γ* < 1 is a ‘future discounting’ factor. Individuals adjust drug intake over time to maximize *R*(*t, u*). We assume that the drug reward is the total subjective reward *R*(*t, u_B_* + *u_D_*(*t*)), and that the mood during withdrawal from the drug is the total subjective reward without drug *R*(*t, u_B_*).

With these definitions, the changes in subjective reward in the addiction process are illustrated schematically in Figure 3C. Initially, drug use, which transiently increases β-endorphin levels, causes an increase in subjective reward (Figure 3C, arrow (i)). This is the initiation phase. After a few weeks, adaptation due to FCD brings β-endorphin and subjective reward back to baseline, despite the increase in drug intake (Figure 3C, arrow (ii)). The circuit therefore produces tolerance to the subjective reward of the drug, leading to escalation of drug intake. Withdrawal from the drug causes a drop in subjective reward (Figure 3C, arrow (iii)) which resolves after several weeks (Figure 3C, arrow (iv)). We conclude that the β-endorphin circuit, in which β-endorphin responds to fold-changes in input, shows the hallmarks of the opponent-process model of addiction.

### FCD circuits are more vulnerable to addiction than other circuits

FCD thus seems to provide the hallmarks for addiction, by providing a fast response and a slow opponent process. We asked whether FCD is special in this regard, or whether other common circuit designs in biology are also prone to addiction. To test this, we compare different prototypical circuits (Figure 3D) in which a drug induces the secretion of a factor which increases subjective reward. We compare the FCD circuit with other circuits that have a slow component with either negative or positive feedback on the secreted factor. We also consider a simple activation topology without a slow component. To make a ‘mathematically controlled comparison’ (Savageau, 1972), we provide all circuits with the same timescale parameters (Methods).

Simulations show that the FCD circuit is unique in the sense that drug reward becomes dissociated from drug intake level and remains constant, and that withdrawal mood drops without bound (Figure 3E). These properties can be shown analytically for all circuits (Methods). Therefore, FCD best captures the properties of the opponent-process model.

To intuitively understand why FCD is especially fragile to addiction, consider the following explanation. At any level of drug intake, the FCD circuit shows a constant (positive) preference for increasing drug intake by 2-fold, and a constant (negative) preference for decreasing drug intake by 2-fold (Figure 3F). This is in contrast with the other circuits, where preference for increasing drug intake, and dislike for decreasing drug intake, both diminish at high levels of drug intake, due to saturation effects. The reason for this is the FCD dynamics presented in Figure 3C, arrow (ii): as drug intake increases, FCD constantly pushes the sensitivity to further intake away from saturation, thus motivating further increases in intake, and demotivating decreases in intake. This may explain the fragility of FCD to addiction, since increasing drug intake does not diminish preference for further increases in drug intake.

### FCD may be advantageous for learning and reward shaping

Finally, we ask what might be the selective advantage of an FCD circuit for reward. We therefore analyze possible advantages of FCD in circuits that control subjective reward.

We focus on the role of reward in the learning of useful behaviors, a major task for cognitive systems. We demonstrate two advantages that FCD confers on learning of behaviors.

Behavior is learned by adjusting actions according to outcomes, such that actions that lead to rewarding outcomes are repeated more often (that is, they are reinforced); actions that lead to aversive outcomes are avoided (Dickinson, 1994; Herrnstein, 1970; Thorndike, 1927). This principle, known as *Thorndike’s law of effect*, is a cornerstone of our understanding of animal learning. It has been mathematically analyzed in the field of reinforcement learning (Dayan and Daw, 2008; Sutton and Barto, 2018).

From the point of view of learning, FCD control of subjective reward is initially puzzling. Algorithms for reinforcement learning, such as the widely-used Q-learning algorithm for optimal action choice, often assume a direct correspondence between the value of an input and its translation into subjective reward. FCD, on the other hand, causes the subjective reward to depend also on the background level of input, breaking the direct correspondence between the input stimulus and its rewarding properties.

Here, we point out that FCD can be crucial for reinforcement learning when rewards span several orders of magnitude. By rescaling inputs according to the background input, FCD allows learning despite large differences in reward values. This FCD feature provides a wide dynamic range over decades of input. An analogous behavior over multiple scales in provided by FCD in systems such as the *E. coli* chemotaxis circuit (Adler and Alon, 2018).

The importance of such scale-invariant sensing for learning has been demonstrated for artificial algorithms such as neural networks (Ioffe and Szegedy, 2015; Santurkar et al., 2018; Sola and Sevilla, 1997). Scale invariance makes gradients more reliable and predictable, reducing the likelihood of vanishing or exploding gradients, and reduces sensitivity to hyperparameters and initialization (Santurkar et al., 2018).

In fact, artificial reinforcement learning algorithms have been shown to benefit from FCD-like processes. Van Hasselt et al. recently added adaptive normalization of the target values of a deep Q-network, which allowed the algorithm to learn to play computer games with varying magnitudes of reward (van Hasselt et al., 2016). This adaptive normalization is similar to FCD, since the rewards were effectively normalized by their background level. This raises the possibility that scale-invariance and FCD may also be important for biological reinforcement learning.

In addition to scale-invariance, FCD has a second important benefit: it provides a natural implementation of a well-established concept in reinforcement learning – *potential-based reward shaping* (Devlin and Kudenko, 2011, 2012; Laud, 2004; Ng et al., 1999; Skinner, 2019). Reward shaping is the addition of auxiliary rewards in order to guide exploration and behavior towards desirable objectives. The motivation behind reward shaping is that ‘real’ rewards, such as access to food or sexual contact, may be sparsely achieved. This sparsity limits the extent to which an agent can learn how to achieve these rewards.

As an example, consider a task presented by Ng. et al (Ng et al., 1999; Randløv and Alstrøm, 1998), where an agent needs to learn to ride a bicycle from point *A* to a distant point *B*. To learn this, the agent receives a reward upon reaching *B*. Since reaching *B* is difficult, the agent gets only sparse feedback for its performance. To address this, the agent is also provided with an auxiliary reward when it approaches B (Randløv and Alstrøm, 1998). However, this leads to a problem, since the agent can now receive reward by simply riding in loops around B, gaining the proximity reward again and again, without ever reaching *B*.

To solve this problem, and provide auxiliary rewards that preserve the learning of optimal behavior, Ng et al. suggested that the agent must be provided with *potential-based auxiliary rewards* (Ng et al., 1999). The intuition is that potential functions have a net effect of zero on any loop. Their net effect depends only on the start and end points. This concept was extended by Devlin and Kudenko (Devlin and Kudenko, 2012) to dynamical potential-based auxiliary rewards. In the standard notation of reinforcement learning, the auxiliary reward for a transition from state *s* at time *t* to state *s*’ at time *t*’ is given by F(s,t,s’, t’) = *γ*Φ(*s*\*, *t*’) – Φ(*s, t*), where *γ* is a discounting factor and Φ is a real-valued potential function. Potential-based reward shaping does not have a problem of loops that can distract the agent. Moreover, potential-based reward shaping preserves the learning of optimal behavior (or *policy invariance*) (Ng et al., 1999).

FCD control of subjective reward can provide a physiological implementation of potential-based reward shaping (Figure 4). The reason for this is that the output of the FCD circuit tracks the logarithmic derivative of its input (Adler and Alon, 2018; Adler et al., 2014; Lang and Sontag, 2016). Thus FCD provides a potential function equal to the log of the input. (see Methods). Such circuits can therefore be thought of as circuits to guide exploration, rather than to learn values.

**Figure 4.**
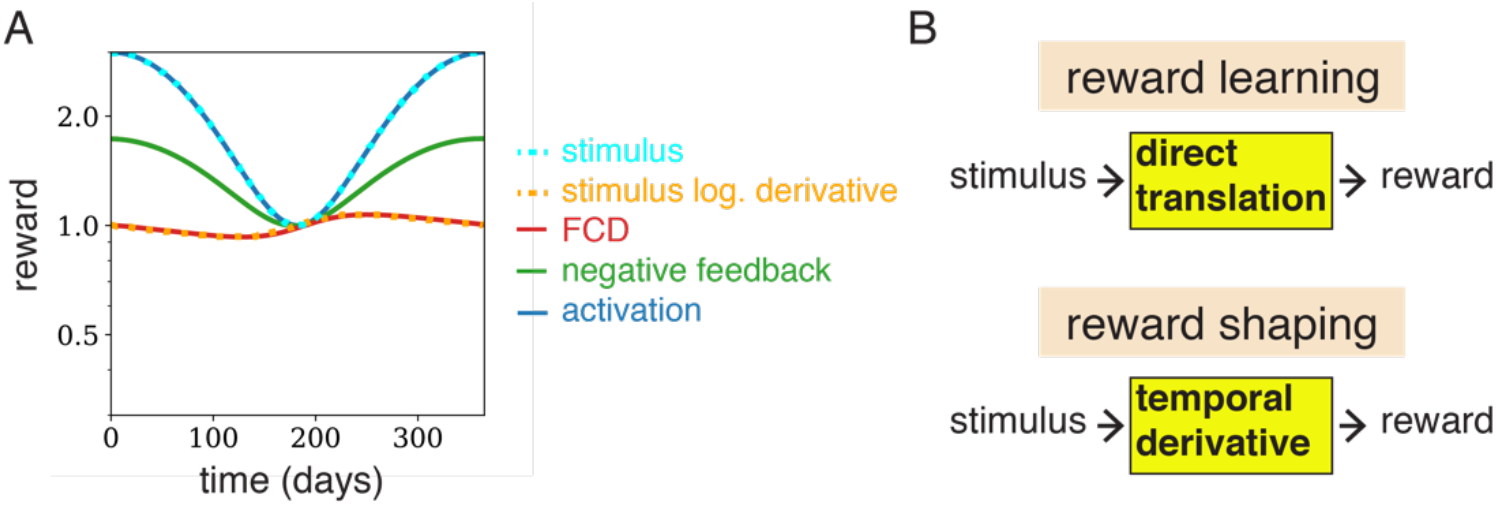
FCD outputs the logarithmic derivative of the input, and therefore provides a physiological implementation of potential-based reward shaping. (A) The negative feedback circuit output (green) and activation circuit output (blue) track the input stimulus (cyan), whereas the FCD circuit output (red) tracks its logarithmic derivative (orange). The input used is 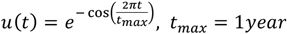. The logarithmic derivative of this stimulus is 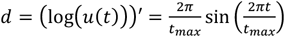, shown scaled and shifted for clarity, 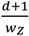 (see methods). (B) Physiological circuits like those discussed in this study convert environmental stimuli to subjective reward. Circuits where the reward corresponds directly to the input stimuli (such as direct activation, negative, or positive feedback) facilitate learning of the input stimuli. On the other hand, circuits like FCD, where the subjective reward tracks the derivative of the input stimuli, facilitate reward shaping, where input stimuli is not directly learned but instead guides exploration towards rewarding behavior.

## Discussion

We propose an opponent process in alcohol addiction based on dynamical changes in the gland masses of the HPA axis. The gland-mass changes due to chronic alcohol intake cause the initial rise in β-endorphin to settle back down to baseline within weeks, and to undershoot for weeks after withdrawal. These dynamics contribute to the physiological basis of the hedonic tolerance and withdrawal stages of addiction. The present opponent process may also apply to other addictive drugs that activate the HPA axis, including cocaine and nicotine (Armario, 2010). This suggests that, in addition to neurotransmitter circuits, endocrine gland mass changes may be potentially important for addiction.

Of particular mechanistic importance in the model is the growth of the adrenal cortex caused by chronic activation of the HPA axis. This growth increases cortisol secretion, which suppresses β-endorphin secretion. Consistent with the model, cortisol levels are about 3-fold higher in people with alcohol abuse disorder during active drinking periods (Stalder et al., 2010); cortisol returns to normal levels after several weeks of abstinence (von Bardeleben et al., 1989; Esel et al., 2001; Marchesi et al., 1997). The model explains why adrenalectomy diminishes alcohol preference in rodents (Fahlke, 2000; Fahlke et al., 1994; Goeders, 2002; Goeders and Guerin, 1996; Hansen et al., 1995; Lamblin and De Witte, 1996). However, the role of HPA gland masses in alcohol abuse disorder or other drug addictions has not, to the best of our knowledge, been directly studied, except for Carsin-Vu et al. (Carsin-Vu et al., 2016) which provide evidence for adrenal enlargement in alcohol abuse disorder.

While our model focused on the role of β-endorphin, it can potentially be generalized to other factors important to alcohol addiction. Any secreted factor that is stimulated by alcohol and inhibited by glucocorticoids, or is secreted as a response to hypothalamic CRH, is predicted by the model to have similar long-term dynamics to β-endorphin (Methods). One potentially relevant factor is dopamine secretion, which is affected by glucocorticoids through interactions with several neural systems (Butts et al., 2011; Piazza and Le Moal, 1996; Piazza et al., 1996). Additional factors include enkephalins which preferentially bind the delta opioid receptor (Froehlich et al., 1991; Marinelli et al., 2005) and are implicated in alcohol addiction (Figure 2B). Future extensions of the present model may include these factors.

The HPA-activating effect of drugs such as alcohol provides reward for mild to moderate stimulation. However, at high levels, limiting mechanisms begin to act. One mechanism is the inhibition of CRH secretion by high levels of cortisol through the low-affinity glucocorticoid receptor GR. Another mechanism is secretion of dynorphins, endogenous opioids which are secreted as a result of prolonged CRH secretion (Nikolarakis et al., 1986). In contrast to β-endorphin, dynorphins have a dysphoric rather than a euphoric effect. This dysphoric effect can presumably prevent over-activation of the HPA axis from being rewarding. Dynorphin secretion was hypothesized to play a role in the dysphoria that follows drug withdrawal (Koob and Volkow, 2016).

The present model infers that increased glucocorticoids during alcohol abuse act to inhibit β-endorphin. This inference is based on experiments done in other contexts, using pharmacological interventions and adrenalectomy. To the best of our knowledge, this inhibition effect has not been directly tested in the context of alcohol abuse disorder. Such experiments will be an important experimental test for the model.

The key mathematical feature that underlies the proposed opponent process is that β-endorphin responds to relative changes in average drug intake, rather than absolute changes, a feature called fold-change detection (FCD). FCD provides the essential hallmarks of the opponent process theory of addiction.

We find that FCD control of reward is therefore especially fragile to addiction. In our model for β-endorphin dynamics, FCD is implemented physiologically by the growth of endocrine glands. FCD in reward pathways can also be implemented in principle by other physiological mechanisms (Adler et al., 2017). For example, adaptation and relative responses to rewards (both hallmarks of FCD) are well documented for midbrain dopamine neurons (Tobler et al., 2005). FCD can occur at the cellular level by biochemical mechanisms such as receptor methylation (Lazova et al., 2011) or by gene-regulation circuits such as incoherent feedforward loops (Goentoro et al., 2009). Regardless of the precise implementation, FCD promotes addiction by maintaining sensitivity to increasing drug intake levels, keeping them away from reward saturation. In this way, wanting the drug is never sated until limiting mechanisms kick in. The relevant timescale for the development of drug tolerance, as well as the affective withdrawal, will depend on the timescale of adaptation in the FCD circuit. While this may be days to weeks for endocrine gland turnover, or possibly for epigenetic modifications, faster circuits (such as transcriptional circuits and protein-protein interactions) may adapt on a timescale of hours, and are therefore less likely to be important for the development of addiction on the timescale of weeks.

It will be interesting to test whether the nicotine receptor system analyzed by Gutkin et al. (Gutkin et al., 2006) also can show FCD (their original model did not generally have FCD). A recent study demonstrated how approximate FCD may be implemented in receptor-based mammalian signaling systems (Lyashenko et al., 2020). In their work, which focused on the pAkt pathway, Lyashenko et al. demonstrated that the adaptation mechanism for FCD can be implemented using receptor endocytosis. Endocytosis occurs generally in neurotransmitter pathways, and may thus be a relevant mechanism for FCD.

In addition to its vulnerability to addiction, FCD control of subjective reward also has selective benefits for learning behaviors that promote fitness. One benefit is to allow rapid learning when reward magnitude spans a large range. This is relevant when the reward baseline level is unknown or fluctuating. A second benefit is a physiological implementation of potential-based reward shaping. This is crucial when the input to the circuit represents a cue or proxy that is correlative with a ‘real’ reward, but does not have value by itself.

Not all behaviorally relevant circuits have FCD: for example, mechanical pain does not adapt to background level (and hence cannot have FCD which entails exact adaptation), whereas pain mediated by interaction with the capsaicin receptor does (Holzer, 1991; Nolano et al., 1999; Winter et al., 1995; Yao and Qin, 2009). It would be fascinating to compare the design principles of different physiological circuits that control subjective reward.

In summary, we propose an opponent process for addiction based on gland-mass changes in the HPA axis. To test this, further experiments are important. Monitoring gland masses using imaging during the stages of the addiction process can help to test this proposal. If gland masses turn out to be important for the week-scale dynamics of addiction, they might serve as relevant targets for intervention. Interventions that suppress gland mass changes at the right time, perhaps using HPA agonists and antagonists, are predicted to interfere with the addiction process and to reduce withdrawal symptoms.

## Materials and Methods

### Model for β-endorphin control by the HPA axis with gland-mass changes

The HPA axis is a cascade of hormones activated by stress inputs. It is initiated by the secretion of CRH from the paraventricular nucleus of the hypothalamus, in response to drugs, as well as to physiological and psychological stresses, whose combined effect is the input denoted u. CRH causes pituitary corticotrophs to secrete ACTH, which, in turn, causes cortisol secretion from the adrenal cortex. CRH also acts as a growth factor for pituitary corticotrophs, and ACTH is a growth factor for the adrenal cortex. To model these dynamics, we used our recent model (Karin et al., 2020), which added to the classic HPA model of Andersen et al. (Andersen et al., 2013) two equations for the changes in gland masses. The non-dimensionalized equations are:

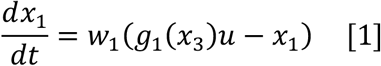

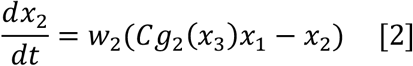

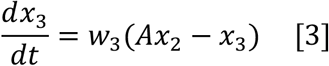

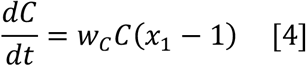

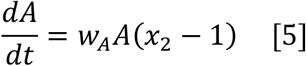

Where *x*_1_, *x*_2_, *x*_3_ are the concentrations of the hormones CRH, ACTH, and cortisol, *C* is corticotroph functional mass, and *A* is the adrenal cortex functional mass. Equations 1–3 for hormone dynamics have a timescale of hours (given by the turnover rates *w*_1_, *w*_2_, *w*_3_). Equations 4–5 have a timescale of weeks due to the gland turnover rates *w_C_* and *w_A_*. All parameters are provided in Table 1. Finally, cortisol has negative feedback on CRH due to both the low affinity receptor GR: 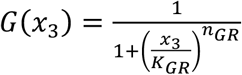 (Andersen et al., 2013; Gupta et al., 2007), and the high affinity receptor MR. Because cortisol saturates MR at physiological levels (Andersen et al., 2013), it is modelled as a Michaelis-Menten function in its saturated regime *M*(*x*_3_) = 1/*x*_3_. Cortisol also has negative feedback on the pituitary due to GR. Therefore *g*_1_(*x*_3_) = *M*(*x*_3_)*G*(*x*_3_), and *g*_2_(*x*_3_) = *G*(*x*_3_).

**Table 1.**
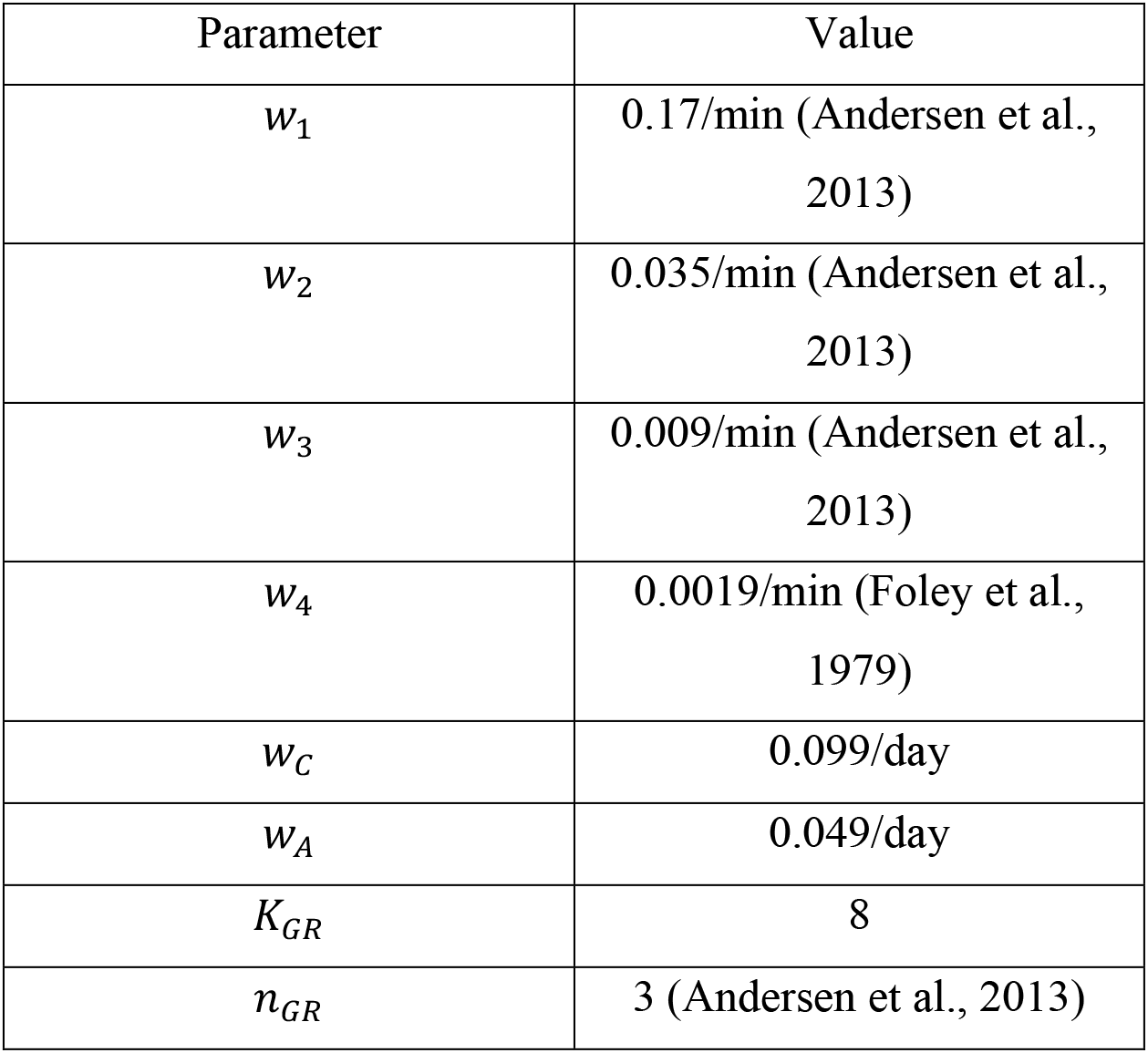
Parameter values for HPA model.

In the present study, we added two new equations for the dynamics of β-endorphin. There are two β-endorphin sources. The first is β-endorphin secreted from pituitary corticotrophs into the bloodstream. This β-endorphin, which we denote by *x*_4_, is co-regulated with ACTH, and therefore is described by the equation:

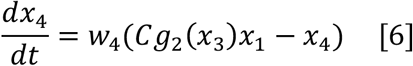

The second case is β-endorphin that is secreted into the cerebrospinal fluid from POMC neurons in the hypothalamus (Bloch et al., 1978; Bloom et al., 1978) in response to CRH, which we denote by *x*_5_:

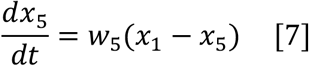

Eq. 7 can also describe β-endorphin secretion from non-corticotroph cells in the pituitary.

For future studies, one may propose a more general equation for a secreted factor *x*_6_ that is stimulated by alcohol and inhibited by cortisol:

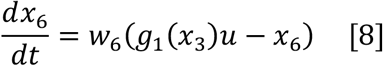

### FCD in β-endorphin dynamics

We now show that *x*_4_, *x*_5_, *x*_6_ all have the FCD property for low to moderate levels of stress input, namely when GR is not appreciably activated, *x*_3_ ≪ *K_GR_*. FCD is defined by dynamics of *x*_4_, *x*_5_ in response to an input *λu*(*t*) that are independent of *λ* > 0, starting from initial conditions at steady-state of all the hormones and glands. To show this, we use normalized variables 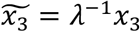 and *Ã* = *λ*^−1^ *A* in Eq. 1–8. Since *x*_3_ ≪ *K_GR_*, the contribution of the GR to the negative feedback is negligible (*G*(*x*_3_) ≈ 1), and thus *g*_1_(*x*_3_) ≈ 1/*x*_3_ and *g*_2_(*x*_3_) ≈ 1. We therefore get:

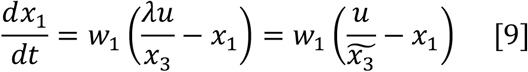

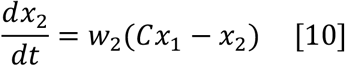

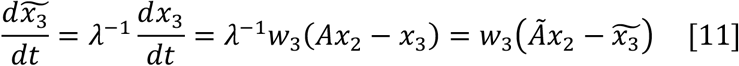

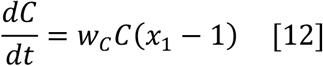

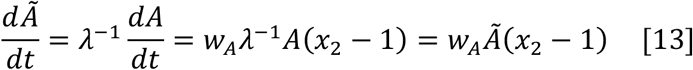

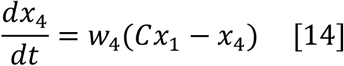

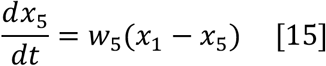

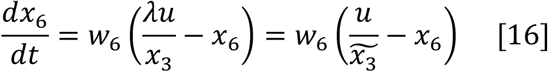

Eq. 8–14 are the same as Eq. 1–17, but without the scale of the input *λ*. In addition, the initial conditions of the hormones are independent of *λ*, since at steady-state *x*_1_ = *x*_2_ = *x*_4_ = *x*_5_ = *x*_6_, and *x*_3_ = *λu* so that 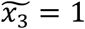. The initial conditions for the gland masses are also independent of *λ*, since at steady state *C* = 1 and *A* = *λu*, so *Ã* = 1. Since both equations and initial conditions are independent of *λ*, the dynamics of *x*_1_, *x*_2_, *x*_4_, *x*_5_, *x*_6_ (as well as *C*) are independent of the scale of the input, *λ*: we obtain the FCD property. Note that this holds only when *x*_3_ ≪ *K_GR_*, otherwise the responses become blunted by the action of GR (the opponent-process property, however, still hold since the negative feedback from cortisol is further strengthened).

### Alternative circuits

Here we provide the equations for the four circuits of Figure 3. In all circuits, we model subjective reward with a generic hill equation: 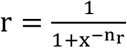, where x is the factor that activates subjective reward (akin to β-endorphin). The hill-coefficient n_r_ determines the sensitivity to changes in x. For the direct-activation circuit, input u directly increases x. For the other circuits, *x* is activated by u and modulated by a slow variable Z. The slow variable *Z*, which provides feedback on the secretion of x, is analogous to the role of the gland functional masses in the HPA axis model (for simplicity of analysis and generality we consider only a single functional mass). The equations for the circuits analyzed in Fig. 3DEF are as follows.

Direct activation circuit:

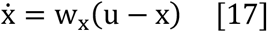

Positive feedback circuit:

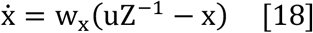

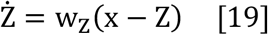

Negative feedback circuit:

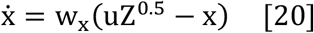

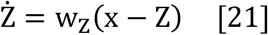

FCD circuit:

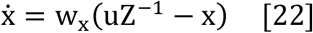

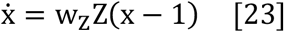

All circuits:

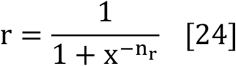

For all circuits w_x_ = 10 min^−1^, w_Z_ = 5day^−1^, n_r_ = 5. The precise values of the powers of Z in the equations do not affect the qualitative results.

### Simulations for FCD responses to steps, pulses, and train of pulses

To generate the simulations in Fig. 3AB, we used the FCD circuit described above, 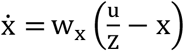, Ż = w_Z_Z(x – 1), with w_x_ = 10 min^−1^, w_Z_ = 5day^−1^, n_r_ = 5. The FCD output plotted is x, which is analogous to β-endorphin in the HPA axis, and Ż is analogous to the adrenal mass in the HPA axis. The input in Fig. 2A consists of two step increases, from u=1 to u=2 at day 20, and from u=2 to u=4 at day 60. In Fig. 3B, the input is given by u(t) = u_B_ + u_D_, where u_B_ = 1 and alcohol intake is 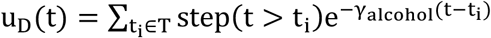 where T = {t_0_,…, t_max_} are the times when alcohol intake occurred. Here we took an alcohol half-life of 6 hours, 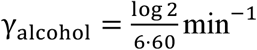. For the upper panel we used T = {1} (in units of days) and for the lower panel T = {1,2,…, 19}. Reward allostasis is achieved because of the slow changes in Z, which cause the average of x to be kept around 1. Too see this, consider an infinite pulse train and two time points t_1_ < t_2_, that both occur before the beginning of a pulse, where t_1_ is large enough so that Z(t_1_) = Z(t_2_). Then: 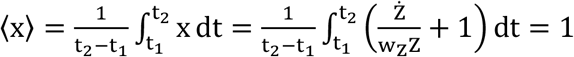.

### Analytical solutions for the effects of escalating drug intake on mood for various circuits

When animals are allowed to self-administrate an addictive drug such as alcohol or cocaine, they gradually escalate their intake over the timescale of days to weeks (Koob, 2013). In this section we analyze analytically how such a gradually increasing input affects the behavior of the various circuit topologies presented in Figure 3. We take a linearly increasing input *u*(*t*) = *w_u_t* where *w_u_* is small compared with *w_s_* and *w_x_*. For all circuits, when *x* is near its steady-state 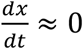, we find *x_ST_* = *uZ^θ^* where *θ* = −1 for negative feedback and FCD circuits and *θ* = 0.5 for the positive feedback circuit. We can substitute this expression in the equation for *Ż*. For FCD we get *Ż* = *w_Z_ Z*(*uZ*^−1^ − 1) = *w_Z_* (*u* – *Z*). Solving this equation gives: 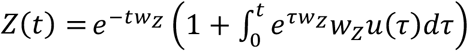. Substituting *u*(*t*) = *w_u_t*, we find

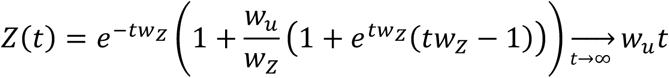

And: 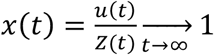. Since drug-related subjective reward *r* is a function of *x*, this suggests that even as drug use escalates, drug-induced reward drops to baseline. We next consider the effect of withdrawal, where drug use drops back to *u* = 1. The new steady-state of *x* (assuming *Z* is still at quasi-steady-state) is 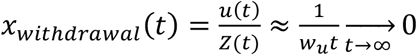. Withdrawal mood drops to zero in the model as drug intake escalates.

For the direct activation circuit, on the other hand, *x* tracks the input *x* = *u*, so drug-induced reward increases with drug intake, and withdrawal mood is the same as the original baseline. Finally, for the other circuits, we get: *Ż* = *w_Z_* (*x* – *Z*) = *w_Z_Z*(*uZ^θ−1^* − 1), which we can solve:

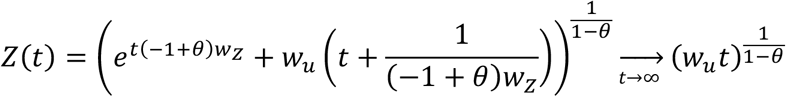

And: 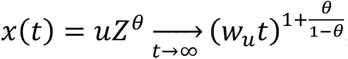, with the withdrawal level of *x* being 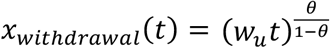. For the negative feedback circuit, where *θ* < 0, drug-induced reward increases sublinearly with the increase in time / drug intake, and withdrawal mood decreases sublinearly with drug intake / time (as the square root for *θ* = −1). Therefore, drug-induced reward remains associated with drug intake, and withdrawal mood, although decreasing, is less sensitive than the FCD circuit to the escalation in drug intake. For the positive feedback circuit, where 0 < *θ* < 1, drug-induced reward increases super-linearly with drug intake (quadratically for *θ* = 0.5), and withdrawal mood also increases with drug intake, in contrast with the observed phenomenology of the addiction process.

### Integral of FCD response

We consider the overall integral of the difference between subjective reward r and its baseline, to a transient perturbation given by u(t), such as the pulse of drug intake shown in Fig. 3A. For simplicity, we assume that the changes in u(t) are small enough so that one can linearize r(t) around x=1, obtaining 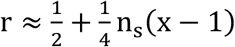. The integral of the response of the FCD circuit is then:

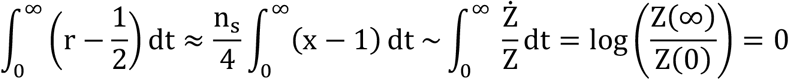

since u(t) changes transiently, Z is the same at t = 0 and at t = ∞. This is not true for the other circuits, and they may therefore have a non-zero integral response to a transient perturbation.

### Model for Reward

Reward optimization was modelled by the standard assumption that individuals try to optimize total discounted subjective reward 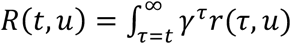 (Dayan and Daw, 2008; Sutton and Barto, 2018). We set *γ* = 0.99, where time is in units of minutes. This value is observed experimentally (McClure et al., 2007), however, the precise value is not important for the conclusions, as long as discounting occurs faster than the timescale of the slow component of the circuit.

### Model for reward shaping

Reward-shaping is defined within the framework of reinforcement learning, where an agent learns to predict the value of its actions (Sutton and Barto, 2018). The decision making of the agent is studied in the context of a Markov Decision Process (MDP). An MDP is a tuple *M* = 〈*S, A, T, R*〉 where *S* is the state space, *A* is the action space, *T*(*s, a, s*′) maps the probability that an action *a* taken at a given state *s* causes a transition to a different state *s*′, and *R*(*s, a, s*′) is the (immediate) reward received for this transition. The goal of optimal learning algorithms such as Q-learning is to find a policy (a map from states to action, Π: *S* → *A*) that maximizes accumulated reward 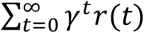, where 0 ≤ *γ* < 1 is a future-discounting factor and *r*(*t*) = *R*(*s_t_, a_t_, s*_*t*+1_) is the immediate reward received at time *t*. This can be generalized to the continuous case, as in (Doya, 2000), by setting the accumulated reward as 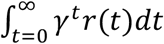.

In order to guide learning, an additional dynamical auxiliary reward function *F*(*s, t, s′, t′*) can be appended, creating a new MDP *M*′ = 〈*S, A, T, R*′〉 where:

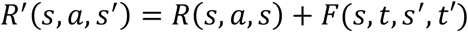

*F*(*s, t, s′, t′*) is the *reward shaping* function that outputs the reward for the transition from state *s* at time *t* to state *s*′ at time *t*′. The MDP *M*’ is guaranteed to preserve the optimal policy of *M* if *F*(*s, t, s′, t′*) can be rewritten as a potential function (Devlin and Kudenko, 2012; Ng et al., 1999):

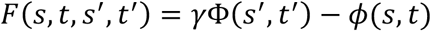

This because the accumulated auxiliary rewards for any sequence of states does not depend on the actions taken:

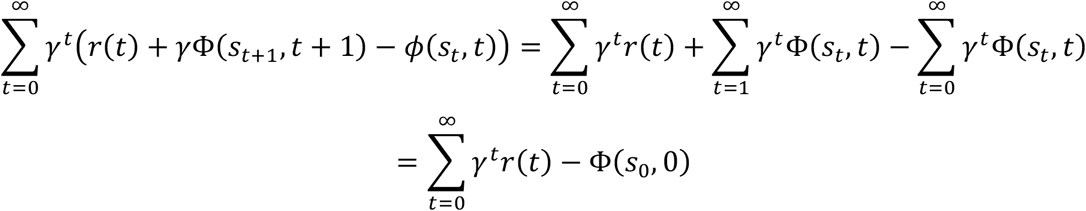

When *γ* ≈ 1, the above holds approximately also when taking:

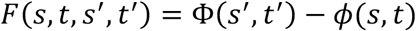

In the continuous case, this is equivalent to setting *F* as the time derivative of a function Φ(s, t):

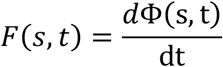

Let us now consider a biological circuit that translates a stimulus that depends on the current state *u*(*s*) to a subjective reward *r*. For circuits like direct activation, negative feedback, or positive feedback, *r* tracks the level of *u*(*s*), as shown in Figure 4. The individual will therefore learn *u*(*s*) through the rewards *r*. On the other hand, for the FCD circuit, *r* tracks the *logarithmic temporal derivative* of *u*(*s*): 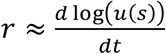. Thus, *r* corresponds to the temporal derivative of a potential function Φ(*s, t*) = log(*u*(*s*)). In this case, the individual will not learn *u*(*s*), but instead *u*(*s*) is used as ‘shaping rewards’ to guide exploration through *r*. FCD can therefore provide a physiological implementation of reward shaping.

### Remark on connection to social attachment

The present model may also relate to aspects of the dynamics of social relations. β-endorphin is thought to play a crucial role in mediating social attachment in primates (Machin and Dunbar, 2011). Social attachment has been suggested to follow distinct stages that are similar in some ways to those of drug addiction, as reviewed by Machin and Dunbar (Machin and Dunbar, 2011). An initial stage of euphoria and attraction develops into tolerance (transition from attraction to attachment). A period of sustained negative affect occurs after separation or social withdrawal. The present model may explain, at least in part, the timescales of these stages since social interactions activate the HPA axis, and therefore β-endorphin dynamics.

The model may also be used to study the combined effects of drug consumption and social interactions. Since both social interactions and alcohol activate the HPA axis, the model predicts that a given serving of alcohol *u_ALC_* will be more rewarding for individuals with a lower baseline of social stimulus *u_SOC_*, since the response is proportional to 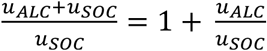. On the other hand, alcohol consumption can also raise β-endorphin levels during social withdrawal, which represents a drop in *u_SOC_*. These observations provide testable predictions that may illuminate social aspects of drug addiction (Heilig et al., 2016).

Changes in HPA gland masses are also well documented in another disorder that is associated with chronic HPA axis activation – major depression. Depressed individuals have increased adrenal mass (Amsterdam et al., 1987; Dorovini-Zis and Zis, 1987; Dumser et al., 1998; Ludescher et al., 2008; Nemeroff et al., 1992; Rubin et al., 1996; Szigethy et al., 1994) which returns to its original size after remission (Rubin et al., 1995). Taken together with the biological and psychological similarities between drug withdrawal and depression (Barr et al., 2002), the present study suggests that gland mass abnormalities may be a common mechanism for affective disorders. If this is indeed the case, interventions aimed at adjusting gland mass size may be beneficial for treating drug addiction and depression.

## Notes

### Competing Interest Statement

The authors have declared no competing interest.

